# SARS-CoV-2 infected cells sprout actin-rich filopodia that facilitate viral invasion

**DOI:** 10.1101/2022.10.19.512957

**Authors:** Yue Zhang, Xiaowei Zhang, Zhongyi Li, Weisong Zhao, Hui Yang, Daijiao Tang, Shuangshuang Zhao, Qian Zhang, Huisheng Liu, Haoyu Li, Bo Li, Pekka Lappalainen, Zongqiang Cui, Yaming Jiu

**Affiliations:** Unit of Cell Biology and Imaging Study of Pathogen Host Interaction, The Center for Microbes, Development and Health, Key Laboratory of Molecular Virology and Immunology, Institut Pasteur of Shanghai, Chinese Academy of Sciences, Shanghai 200031, China; University of Chinese Academy of Sciences, Yuquan Road No. 19(A), Shijingshan District, Beijing, China; State Key Laboratory of Virology, Wuhan Institute of Virology, Center for Biosafety Mega-Science, Chinese Academy of Sciences, Wuhan, China; Institute of Biomechanics and Medical Engineering, Applied Mechanics Laboratory, Department of Engineering Mechanics, Tsinghua University, Beijing 100084, China; Advanced Microscopy and Instrumentation Research Center, School of Instrumentation Science and Engineering, Harbin Institute of Technology, Harbin, China; Guangzhou Laboratory, Guangzhou, Guangdong, China; Institute of Biotechnology and Helsinki Institute of Life Science, University of Helsinki, P.O. Box 56, 00014, Helsinki, Finland

**Keywords:** SARS-CoV-2, filopodia, viral invasion, actin cytoskeleton, anti-viral therapy

## Abstract

Emerging COVID-19 pandemic caused by severe acute respiratory syndrome coronavirus 2 (SARS-CoV-2) poses a great threat to human health and economics. Although SARS-CoV-2 entry mechanism has been explored, little is known about how SARS-CoV-2 regulates the host cell remodeling to facilitate virus invasion process. Here we unveil that SARS-CoV-2 boosts and repurposes filopodia for entry to the target cells. Using SARS-CoV-2 virus-like particle (VLP), real-time live-cell imaging and simulation of active gel model, we reveal that VLP-induced Cdc42 activation leads to the formation of filopodia, which reinforce the viral entry to host cells. By single-particle tracking and sparse deconvolution algorithm, we uncover that VLP particles utilize filopodia to reach the entry site in two patterns, ‘surfing’ and ‘grabbing’, which are more efficient and faster than entry via flat plasma membrane regions. Furthermore, the entry process via filopodia is dependent on the actin cytoskeleton and actin-associated proteins fascin, formin, and Arp2/3. Importantly, either inhibition the actin cross-linking protein fascin or the active level of Cdc42 could significantly hinders both the VLP and the authentic SARS-CoV-2 entry. Together, our results highlight that the spatial-temporal regulation of the actin cytoskeleton by SARS-CoV-2 infection makes filopodia as a ‘highway’ for virus entry, which emerges as an antiviral target.

**Significance Statement:** Revealing the mechanism of SARS-CoV-2 invasion is of great significance to explain its high pathogenic and rapid transmission in the world. We discovered a previously unknown route of SARS-CoV-2 entry. SARS-CoV-2 virus-like particles boost cellular filopodia formation by activating Cdc42. Using state-of-art-technology, we spatial-temporally described how virus utilize filopodia to enter the target cell in two modes: ‘surfing’ and ‘grabbing’. Filopodia can directly transport the virus to endocytic hot spots to avoid the virus from disorderly searching on the plasma membrane. Our study complements current knowledge of SARS-CoV-2 that filopodia and its components not only play an important role in virus release and cell-cell transmission, but also in the entry process, and provides several potential therapeutic targets for SARS-CoV-2.

**Highlights:**

- SARS-CoV-2 VLP infection promotes filopodia formation by activating Cdc42
- SARS-CoV-2 VLP utilizes filopodia to enter target cell via two modes, ‘surfing’ and ‘grabbing’
- Filopodia disruption compromises the invasion of both VLP and authentic SARS-CoV-2

## Introduction

The COVID-19 pandemic caused by severe acute respiratory syndrome coronavirus 2 (SARS-CoV-2) has resulted in more than 600 million infections and 6.4 million deaths worldwide since its emergence in the late 2019, which seriously affects global health and economy (Onyeaka et al. 2021). Despite the rapid development of relevant therapeutic studies and vaccines, the extensively spread of emerging variants of SARS-CoV-2 emphasizes the urgency of developing effective broad-spectrum pharmacotherapeutic interventions. Therefore, it is important to enrich the understanding of the very first step upon SARS-CoV-2 infection, invasion mechanisms, especially from the perspective of host cell morphological deformations, which will provide critical structural basis for viral entry, but has not been raised more attention. SARS-CoV-2 virion comprises a positive-sense single-strand RNA decorated by the nucleocapsid protein (NP), and viral membrane decorated by the structural proteins envelop (E), membrane (M) and spike (S). After binding to the angiotensin-converting enzyme 2 (ACE2) receptor via receptor-binding domain (RBD) of S protein, SARS-CoV-2 completes invasion through two distinct pathways: a fast route via plasma membrane fusion and a relatively slow route via endocytosis (Jackson et al. 2022). The fusion process can be activated by transmembrane protease serine 2 (TMPRSS2), if the host cells express it, on the plasma membrane, in which case the virus can rapidly enter the cell within 10 min in a pH-independent manner (Koch et al. 2021). On the other hand, when targeted cells lack of TMPRSS2 expression, virus-ACE2 complex is internalized via clathrin-mediated endocytosis which is usually taking 40 ~ 60 min (Bayati et al. 2021, Koch et al. 2021). Although SARS-CoV-2 entry process has been investigated since the beginning of the outbreak, little is known about the spatial- and tempo-dynamics of viral particle trafficking and virus-cell interactions during entry. Filopodia are thin, finger-like plasma membrane protrusions of up to several microns in length (0.1 ~ 0.3 μm), that are involved in biological functions such as cell adhesion, cell migration and environmental probing (Mattila and Lappalainen 2008). Filopodia contain bundles of long parallel actin filaments and exhibit highly dynamic protrusions and retractions, which are determined by the balance between actin polymerization at the tip and retrograde flow at the base of filopodium (Mallavarapu and Mitchison 1999, Mattila and Lappalainen 2008). Many proteins are involved in regulating filopodia formation, such as actin cross-linking protein fascin, parallel actin nucleating protein formin, and branched actin nucleating protein Arp2/3 (Mellor 2010, Aliyu et al. 2021). Recent studies reveal that larger number of elongated filopodia, microvilli, actin bundles, and stress fibers induced by SARS-CoV-2 infection are important for virus egress and/or cell-to-cell spread (Bouhaddou et al. 2020, Barreto-Vieira et al. 2021, Nijenhuis et al. 2021). However, the morphological dynamics and the biological roles of filopodia during the invasion of SARS-CoV-2 remain poorly characterized.

Here, we constructed florescence tagged virus-like particle (VLP) of SARS-CoV-2, and utilized the single-particle tracking to visualize the dynamic invasion process of SARS-CoV-2 in VeroE6 cells. We uncovered that SARS-CoV-2 VLPs induced robust formation of filopodia-like structures, and hijacked the inherent host filopodia machinery to transport viral particles into the cell. Real-time imaging unambiguously reveals two characteristics of filopodia-based SARS-CoV-2 internalization, ‘viral surfing’ and ‘viral grabbing’. By carrying out series drugs perturbation experiments, we confirmed that VLP internalization into host cells proceed with requirement of filopodia components, fascin, formin, Arp2/3, and Cdc42 GTPase activity. Concurrently, either disruption of fascin or inhibition of Cdc42 activity prevents the invasion efficiency of authentic SARS-CoV-2. Together, our work provides direct evidence on the involvement of filopodia during SARS-CoV-2 invasion, which is crucial for understanding the initial steps of SARS-CoV-2 infection, and provides profound insights into the development of novel and broad-spectrum antiviral interventions.

## Results

### SARS-CoV-2 VLP infection enhances the filopodia formation by enhancing Cdc42 GTPase activity in host cells

To investigate the dynamic process of how SARS-CoV-2 enters host cells, we applied live-cell imaging using fluorescently labeled virus-like particle (VLP). S (spike)-EGFP gene was co-transfected with the E (envelope), M (membrane), and NP (nucleocapsid) genes into HEK293T cells, to produce EGFP-labeled SARS-CoV-2 VLPs (VLPs-EGFP) (Fig S1A). Purified VLP samples were analyzed with transmission electronic microscope (TEM) and revealed spherical particles of approximately 100 nm in size with a pronounced crown or ‘corona’, with no difference in morphology between VLPs-EGFP and wild-type VLPs (Fig S1B). In an immunofluorescence assay, VLP samples immunostained with specific antibodies against SARS-CoV-2 NP and S protein, respectively. Almost all the NP or S signals were colocalized with EGFP fusion protein, confirming the successful assembly of SARS-CoV-2 VLPs-EGFP (Fig S1C). VeroE6 cells incubated with VLPs-EGFP showed clear fluorescent signal in the cytoplasm (Fig S1E), indicating that EGFP labeling did not impair the entry of viral particles. What’s more, we constructed VeroE6-ACE2 cells (VeroE6 wild-type cell line stably overexpressing exogenous ACE2) by lentivirus system to increase the infection efficiency of VLPs-EGFP, verifying by western blot and imaging method (Fig S1D-E). We then tested the enter dynamics, and measured the number of VLPs-EGFP particles in infected cells at 3 hpi to ensure most of VLPs have entered the cell. Furthermore, live-cell imaging was started to record from 0.5 hpi (starting time point for the exponential rising period of VLP entry) onwards to capture the process in which VLPs enters the host cells (Fig S1F).

To visualize and follow the dynamic morphological changes of infected cells, fluorescently tagged wheat germ agglutinin (WGA) was used to stain plasma membrane and track protrusive structures (Twarock et al. 2010). Cells seeded at a low confluency (avoid cell-cell contact) were infected with VLPs-EGFP for 30 min at 37°C, and observed continuously by high-resolution spinning disc confocal microcopy for the following 2 h. Intriguingly, we found that VLP infection induced robust formation of filopodia-like structures on the host cell surface (Fig 1A, S1G, Video S1-2). To gain more insights in the morphodynamic characteristics of filopodia, we quantified a series of parameters, including the maximum length to represent filopodia shape, the number and persistent time to represent filopodia dynamics, the angle relative to plasma membrane, the protruding speed, and the retracting speed to represent filopodia movement (Fig 1B-G). Compared to un-infected cells, the number of longer filopodia was greatly increased, and the movement speed of filopodia was significantly accelerated, without affecting the entire persistent duration of filopodia in VLP-infected cells (Fig 1B-G), revealing that VLP infection boosted the formation and dynamics of elongated filopodia. Nevertheless, VLP infection has no influence on protrusion angle of filopodia (Fig 1E), suggesting that the overall sensing orientation of filopodia is not specifically changed upon infection.

**Figure 1.**
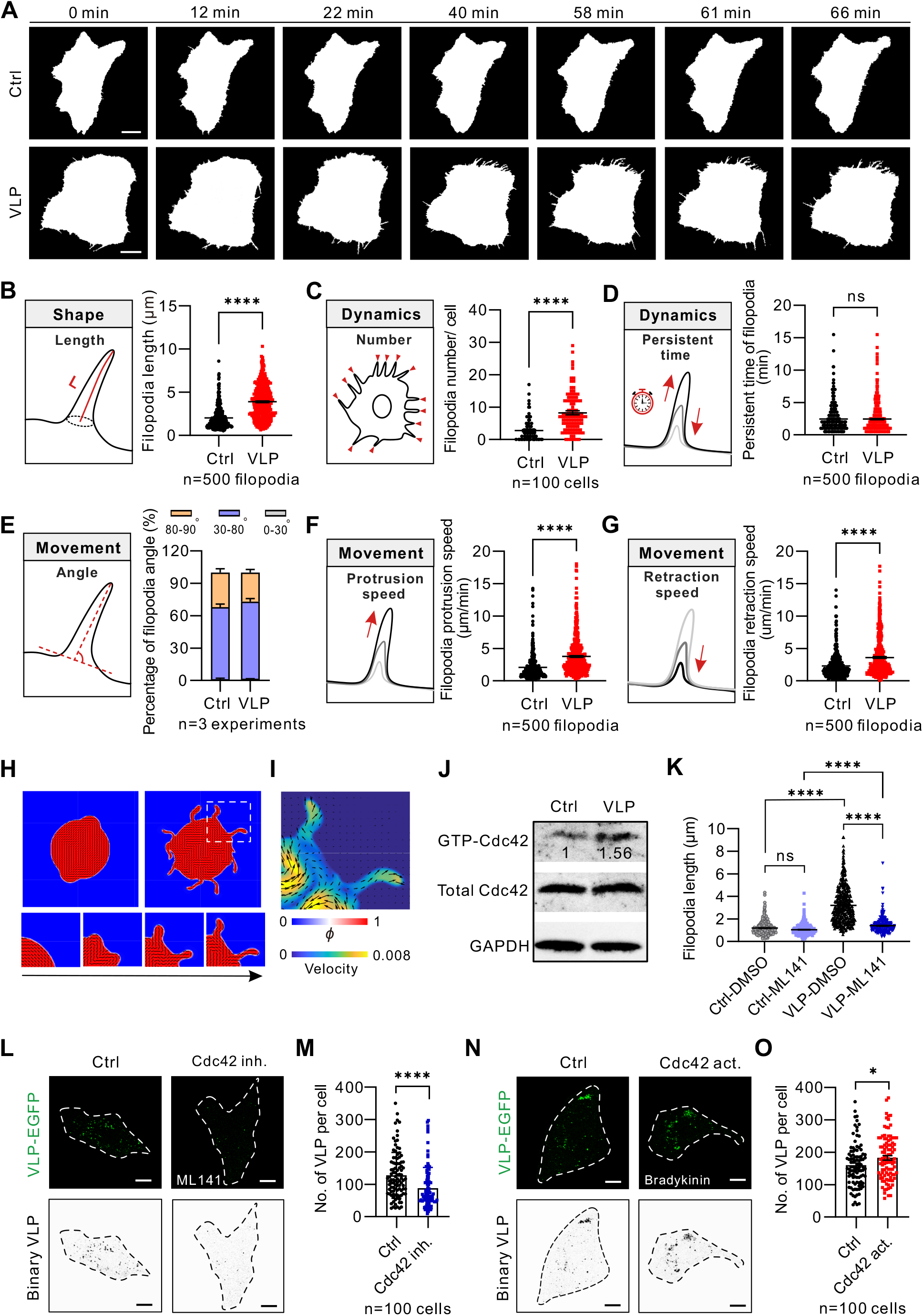
SARS-CoV-2 VLP infection induces filopodia formation by enhancing Cdc42 GTPase activity in host cells. (**A**) Representative binary images from 2 h time-lapse videos of untreated (Ctrl) and VLP infected Vero-E6-ACE2 cells. Scale bar 10 μm. (**B**) Quantification of filopodia length in Ctrl and VLP infected cells. (**C**) Quantification of the numbers of filopodia per Ctrl or VLP infected cells. n=100 cells from three independent experiments. (**D**) Quantification of the persistent time of filopodia in Ctrl and VLP infected cells. (**E**) Quantification of protrusive angles of filopodia relative to plasma membrane in Ctrl and VLP infected cells. The data were from three independent experiments. (**F**) Quantification of the protrusion speed of filopodia in Ctrl and VLP infected cells. (**G**) Quantification of the retraction speed of filopodia in Ctrl and VLP infected cells. n=500 filopodia from three independent experiments in (B), (D), (F) and (G). (**H**) The formation and evolution of filopodia were captured by the active gel model. The left-upper and right-upper panels show the morphology of a cell at low activity (ζ=0.005) and high activity (ζ=0.02), respectively. The lower panels show the dynamical process of filopodial elongation from the white square area in the upper panel. Regions with higher and lower *ϕ* are shown in red and blue, respectively. (**I**) Local direction and velocity of actin from the white square area of (H). The arrows indicate the actin direction, and the short lines indicate the velocity. (**J**) VLP-induced activation of Cdc42 in VeroE6-ACE2 cells. Numbers in the blots indicated the levels of active GTP-Cdc42 normalized to total Cdc42 level and Ctrl cells. (**K**) Quantification of filopodia length in DMSO or ML141 treated Ctrl and VLP infected cells. The data are represented as means ± SEM of 500 filopodia from three independent experiments. (**L, N**) Representative images of the VLP particles entered VeroE6-ACE2 cells under untreated (Ctrl), cdc42 inhibitor ML141 (L) and cdc42 activator bradykinin (N) treated cells. VeroE6-ACE2 cells were treated with 500 ng/mL bradykinin or 50 μM ML141 for 5 h, infected with 0.15 mg VLPs for 3 h at 37°C and fixed with 4% PFA for 15 min at RT. White dotted lines indicate cell outlines that defined by bright field. Binary images shown at the bottom of the picture is for clearer presentation. (**M, O**) Quantification of the number of VLPs shown in (L) and (N). The data are represented as means ± SEM of 100 infected cells from three independent experiments. ns (no significant difference) *P* > 0.05, \**P* ≤ 0.05 and \*\*\*\**P* ≤ 0.0001 (unpaired *t*-test).

To elucidate the physical mechanism underlying the formation and growth of filopodia in host cells subjected to SARS-CoV-2 VLP infection, we proposed an active gel model, in which we used a scalar field □ to describe the shape of the cell and introduced an order tensor **Q**=2*S*(**nn**-**I**/2) which is coupled to □, to describe actin cytoskeleton. We hypothesized intuitively that filopodial protrusion or contraction was driven by active interaction forces between actin filaments in host cells, described by the active stress **σ**_act_ = -*ζ***Q** (Giomi et al. 2013, Marchetti et al. 2013, Julicher et al. 2018, Li et al. 2020). In the equation, activity coefficient ζ>0 corresponds to the extensile interaction and ζ<0 corresponds to the contractile interaction, both of which were modulated by the external stimuli from SARS-CoV-2 VLPs. The active stress drives a flow **v**, which in turn mediates the shape of the host cell. The formation and evolution of filopodia can thus be captured by numerically solving the proposed active gel model. We first simulated the morphology of host cell without SARS-CoV-2 VLP infection by setting a low activity, where cells displayed stable, relatively rounded configuration and did not generate spiky filopodia (Fig 1H), recapitulating the control group in our experiment (Fig 1A). Upon SARS-CoV-2 VLP infection, the activity of cells was enhanced (e.g., ζ=0.02), which can spontaneously form a number of slender filopodia growing radially on the cell surfaces (Fig. 1H), substantiating our experimental observation (Fig 1A-G). We further illustrated the local actin cytoskeletal structure and the velocity field of filopodia (Fig 1I). It appears that the actin flows driven by active stress may readily break the symmetry of host cells, initiating the protrusion of finger-like filopodia. Subsequently, actin cytoskeleton self-organized and flowed along the filopodia, further driving the elongation of the filopodia (Fig 1H, 1I).

Apart from physical perspective, Rho GTPase Cdc42 has been generally acknowledged as the biochemical mechanism closely linked to the dynamic assembling of actin cytoskeletal filopodia (Ridley 2006). We subsequently evaluated the active level of Cdc42 upon infection by GTPase pull down assay. Meanwhile, Cdc42 activator bradykinin and inhibitor ML141 were utilized to further investigate the mechanism of VLP induced filopodia formation. The results showed that VLP infection indeed enhanced the active level of Cdc42 (Fig 1J). ML141 treatment significantly shortens the filopodia length that boosted by VLP infection, but did not affect un-infected cells (Fig 1K). Reciprocally, inhibition of Cdc42 significantly reduced the number of entered VLPs, while activation of Cdc42 reinforced the number of entered VLPs (Fig 1L-O). Together, these results confirm that VLP infection enhanced the formation and dynamics of filopodia in host cells by activating Cdc42.

### SARS-CoV-2 VLPs enter host cells via filopodia in two patterns

To investigate whether SARS-CoV-2 VLPs associate with filopodia to facilitate their invasion process, single-particle tracking was subsequently performed. In order to optimize the imaging data to unambiguously visualize both thin filopodia and small VLP particles in real-time, we first used a percentile-based normalization to reduce the large intensity differences between the dense cytoplasmic signal and the isolated filopodia signal. Then, considering the weak fluorescence signal of the VLPs-EGFP, we used the recent developed algorithm, sparse deconvolution, to eliminate the read-out noise and enhance the contrast of the low signal-to-noise ratio (Zhao et al. 2022) (Fig 2A).

**Figure 2.**
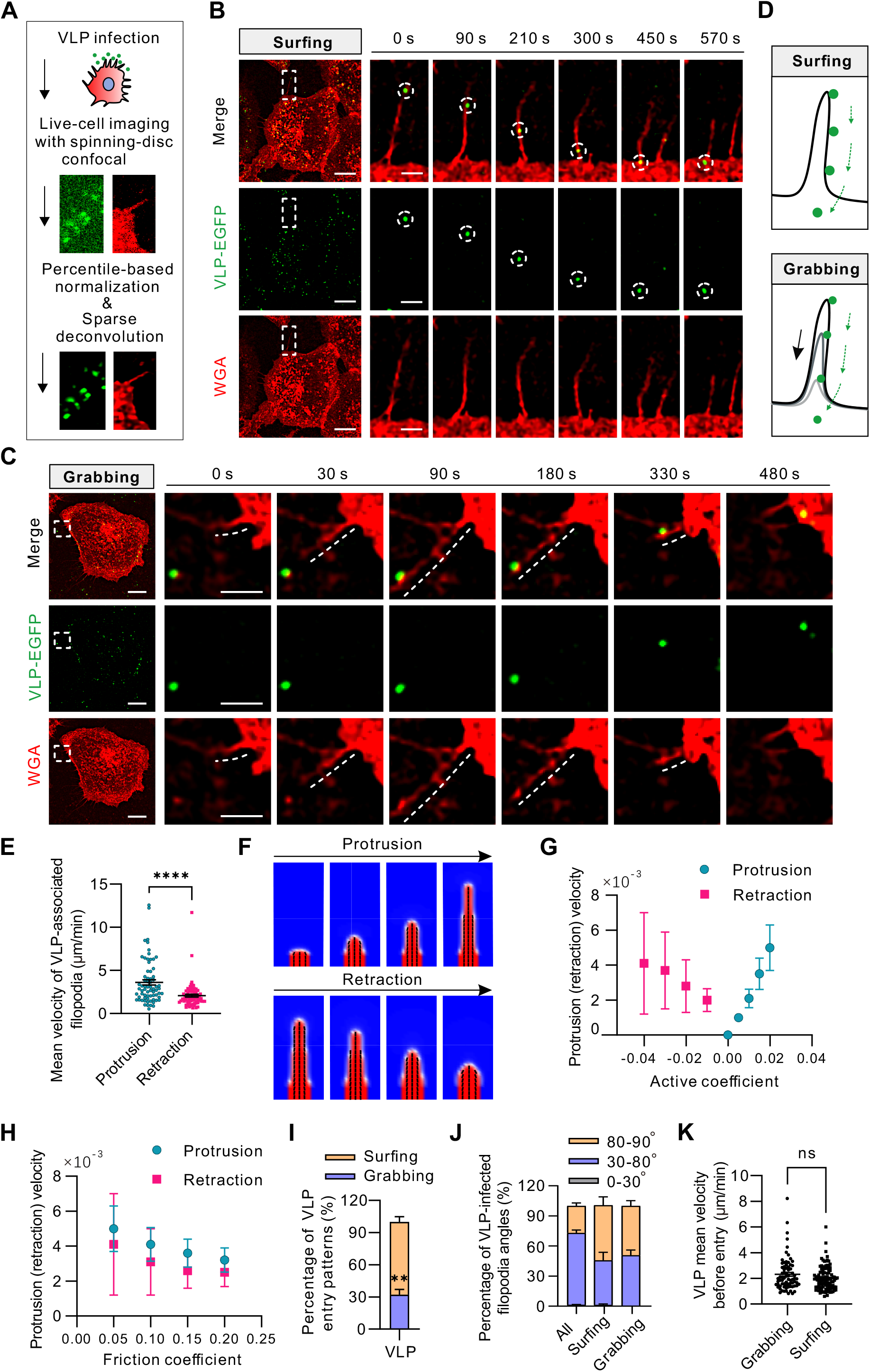
Two invasion patterns of SARS-CoV-2 VLP via filopodia. (**A**) Work flow of single-particle tracking and data analysis. (**B**) A representative VLP particle surfs on VeroE6-ACE2 cell. White rectangles indicate the magnified area shown in the corresponding right panel. The dotted circles indicated the positions of the VLP foci. Cell outlines were stained by WGA. Scale bar 10 μm in the cell images and 2 μm in the magnified images. (**C**) Representative example of filopodial protrusion grabs one VLP particle. White squares indicate the magnified area of the representative images shown in the corresponding right panel. The dotted white lines indicated the filopodia shape stained by WGA. Scale bar 10 μm in the cell images and 3 μm in the magnified images. (**D**) Schematic model of two patterns of VLP invading cells via filopodia, surfing and grabbing. (**E**) Quantification of the mean velocity of VLP-bound filopodia. Each point represents one filopodia. (**F**) Simulation of the evolution dynamics of a single filopodium. Filopodial elongation and retraction are under the extensile and contractile activity, respectively. (**G**) Filopodial speed versus the activity coefficient in (F). **(H)** Filopodial speed versus the friction between the cell and the underlying substrate in (F). (**I**) Quantification of the percentage of VLP entry patterns via filopodia. (**J**) Quantification of the percentage of filopodia angles in VLP infected VeroE6-ACE2 cells. ‘All’ indicates the overall filopodia in infected cells. ‘Grabbing’ indicates the filopodia that grab VLPs. ‘Surfing’ indicates the filopodia that VLPs surf on it. (**K**) Quantification of the mean velocity of VLP before entry. Each point represents one VLP particle. VeroE6-ACE2 cells were infected with 0.15 mg VLP for 0.5 h at 37°C, stained with WGA for 10 min and imaged for 1.5 h. The data are represented as means ± SEM from three independent experiments in (E, I-K). ns (no significant difference), *P* > 0.05, ** *P* ≤ 0.01 and \*\*\*\**P* ≤ 0.0001 (unpaired *t*-test).

By monitoring the motion of individual viral particle through the early stage of infection, we observed that VLPs not only attached to filopodia but also underwent directed rearward movement surfing toward the cell body (Fig 2B, Video S3). In addition to surfing, we found another unique spatial-focusing process that a VLP particle was grabbed by filopodia from its initial random landing position back into the cell body (Fig 2C, Video S4). In other words, there seems an active searching process governed by filopodia from the time it starts to protrude out of plasma membrane. Referred to previous studies, we termed these two filopodia-dependent patterns of VLP entering as ‘surfing’ and ‘grabbing’, respectively (Fig 2D). Of note, the retraction speed of filopodia was significantly slowed down when loaded with VLP, compared to the speed of unloaded filopodia (Fig 2E).

To elucidate the physical mechanism underlying the ‘grabbing’ mode, we simulated the evolution of a single filopodium, which indicated that the high extensile activity induced by SARS-CoV-2 VLP infection speeds up the elongation of filopodium (Fig 2F-G). Moreover, simulation results predicted that the speed of filopodial elongation or retraction depended on the interfacial interaction between the filopodium and the underlying substrate. The increasing friction resistance at the substrate surface slowed down the actin flows and, thereby, reduced the speed of filopodia (Fig. 2H). Inferably, the VLP loading enhance the friction resisting filopodial movement, because of either increased pressure arising from VLP gravity or repeatedly surface trapping of VLP in roughness troughs, which aligns with our experiment that the retraction speed of filopodia became remarkably slow when loaded with VLP (Fig. 2E). Comparing these two filopodia-dependent entry patterns, surfing events appeared more frequently (~70%) than grabbing based on our observation (Fig 2I). Of note, filopodia have already been established before binding of VLP particles in ‘surfing’ mode, and normally remain its structure without retracting back to the cell body (Fig 2B). Whereas the angle of success-VLP-grabbing filopodia is consistently tend to almost 90 degrees, indicating that these filopodia may protrude purposefully to uptake the VLP particle with the shortest reaching distance (Fig 2C,2J). Further quantification analysis revealed that both surfing and grabbing enabled VLP to enter target cells within 10 min. The mean velocity and entry time of VLP have no significant difference between these two patterns (Fig 2K). The slower retraction rate of filopodia was highly accordant with the mean speed at which VLP particle enters the host cell (Fig 2E,2K).

Apart from filopodia-dependent mode, single-particle tracking also revealed that SARS-CoV-2 VLPs enter host cells via plasma membrane (Fig 3A-B). Interestingly, some VLPs can enter the cell directly and being transported inwardly within 10 min, while some VLPs continually move laterally on the plasma membrane and take substantially longer time for entry (Fig 3A-B, Video S5-6). We classified above plasma membrane dependent patterns as ‘fast mode’ and ‘slow mode’ based on the duration of VLP entry (Fig 3C). The quantifications demonstrated that slow mode appeared more frequently than fast mode (Fig 3D), and the taken time in slow mode is consistent with that reported in the previous literature (Bayati et al. 2021, Koch et al. 2021), about 20 ~ 60 min (Fig 3E). Moreover, it is worth noting that compared to the slow mode of plasma membrane dependent entry, all VLP particles utilizing filopodia take substantial less time to enter cells (Fig 3E-F). Combing results of VLP entry patterns, it is reasonable to presume that SARS-CoV-2 VLPs utilize cellular filopodia to act as a ‘highway’ to enter the target host cells, which avoids VLPs from the process of searching a suitable entry site on the cell membrane, therefore, enhances the entry efficiency of VLPs.

**Figure 3.**
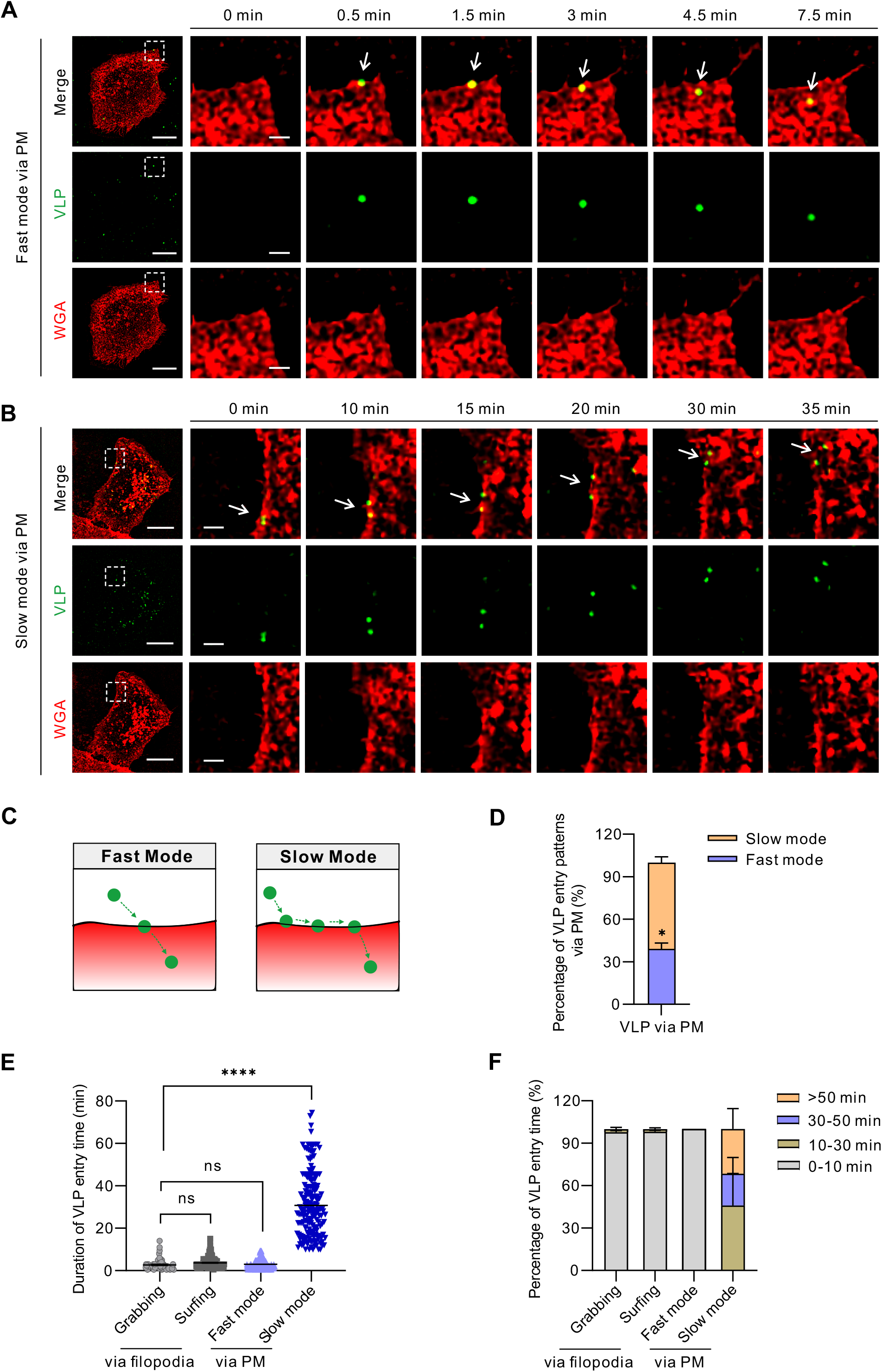
Two entry patterns of SARS-CoV-2 VLP via plasma membrane. (**A, B**) Representative real-time images of VLPs entry into target cell via plasma membrane in fast mode (A) and slow mode (B). White dotted rectangles indicate the magnified area shown in the corresponding right panel. White arrows point the position of VLP foci. Scale bar 15 μm in the cell images and 2 μm in the magnified images. (**C**) Schematic model of two patterns of VLP invading cells via plasma membrane, fast mode and slow mode. (**D**) Quantification of the percentage of VLP entry patterns via plasma membrane. (**E**) Quantification of the time duration for VLP entry via filopodia and plasma membrane, respectively. (**F**) Classification of the time duration for VLP entry in (E). The data are represented as means ± SEM from three independent experiments in (D-F). ns (no significant difference), *P* > 0.05, * *P*≤ 0.05 and \*\*\*\**P* ≤ 0.0001 (unpaired *t*-test).

### Actin associated filopodia components are essential for the entry of SARS-CoV-2 VLP

There are a key set of proteins that have been reported to be involved in filopodia formation (Fig 4A) (Mattila and Lappalainen 2008). To further dissect the role of filopodia in SARS-CoV-2 VLP infection, a series of essential components of filopodia structure were disrupted by chemical inhibitors (Fig 4B). First, we confirmed the optimal dosage and administration duration of all drugs by observing their effects on actin filaments and filipodia characterization. Even treatment with very low concentration, latrunculin B (0.1 μM for 40 min) exhibits cytotoxicity effect to cells (Fig S2A), disrupts actin filaments organization, significantly changes the cell morphology, and depletes filopodia from the cell surface (Fig 4C). Treatment with NP-G2-044 (50 μM for 2 h), CK666 (100 μM for 2 h), blebbistatin (17 μM for 1 h) and SMIFH2 (15 μM for 1 h) significantly reduced the length, protrusion and retraction speed of filopodia, without affecting protrusion angle and severe cytotoxicity, but increased the number of filopodia per cell (Fig 4E-I, S2B-F). Specifically, when treated with myosin II inhibitor blebbistatin, cells formed less stress fibers and acquired a more dendritic morphology (Fig 4C).

**Figure 4.**
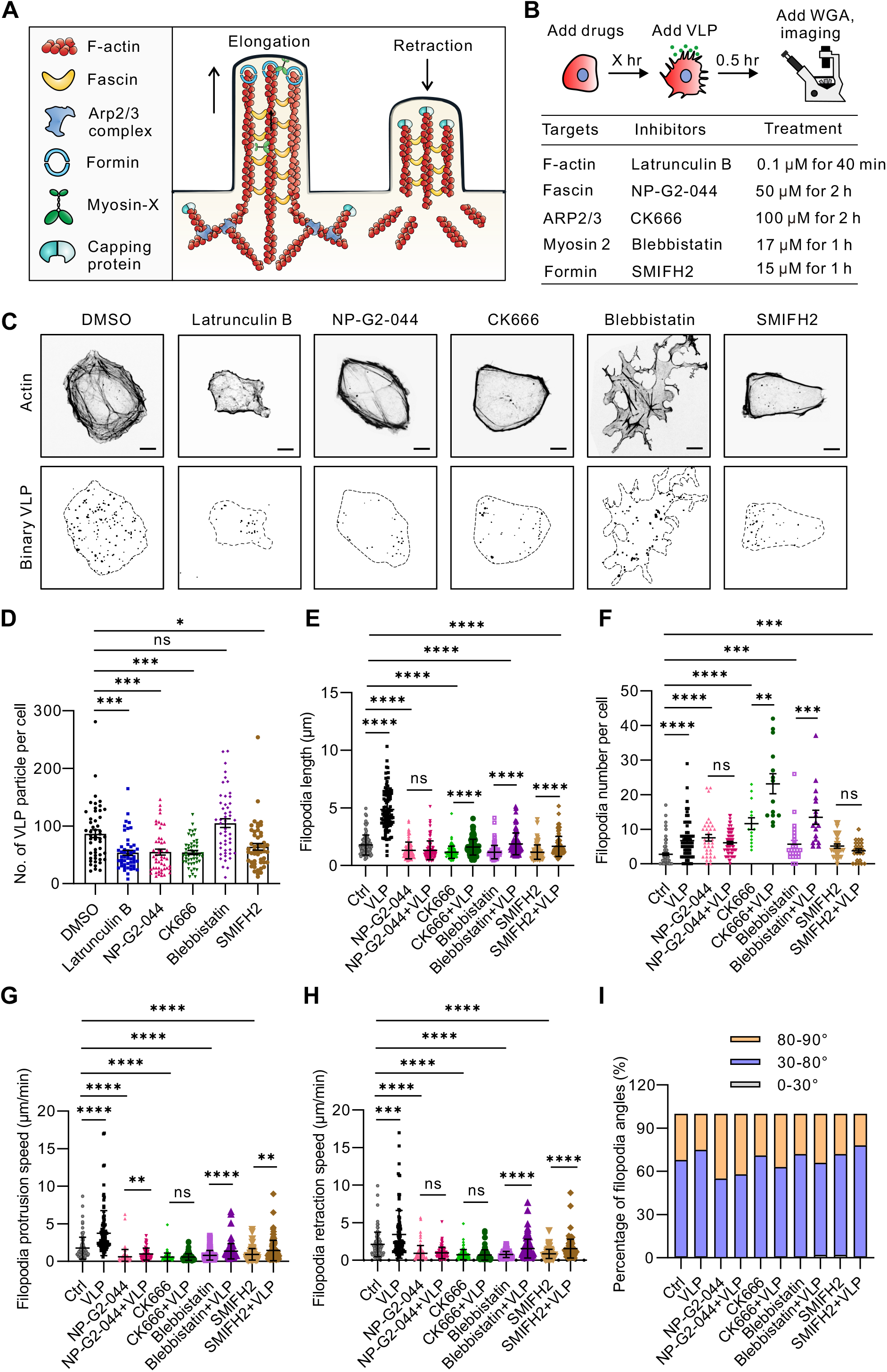
Disruption of filopodia inhibits SARS-CoV-2 VLP invasion. (**A**) Schematic model describes functions of key proteins contributing to filopodia formation. (**B**) Workflow of drugs treatment during live-cell imaging. (**C**) Representative images of drugs treated cells in Ctrl and VLP-infected condition. Scale bar 10 μm. (**D**) Quantification of the numbers of VLP per cell in (C) upon VLP infection at 3 hpi. The data are represented as means ± SEM of 50 infected cells from three independent experiments. (**E-I**) Quantification of filopodia length (E), filopodia number per cell (F), filopodia protrusion speed (G), filopodia retraction speed (H), filopodia angle (I) in untreated (Ctrl) and drug treated cells upon VLP infection. The data are represented Means ± SEM from three independent experiments. ns *P* > 0.05, \**P* ≤ 0.05, \*\**P* ≤ 0.01, \*\*\**P* ≤ 0.001 and \*\*\*\**P* ≤ 0.0001 (unpaired *t*-test).

To compare the entry efficiency of VLPs, VeroE6-ACE2 cells were pretreated with these drugs, and subsequently infected with VLPs in the presence of drugs for another 3 h at 37°C. The inhibitory effect of drugs was proved by staining actin network by phalloidin upon fixation. We quantified the number of intracellular VLP particles by Imaris Spots configuration, and found that treatment with NP-G2-044, CK666, latrunculin B and SMIFH2 significantly reduced the number of VLP particles within host cells, except for blebbistatin (Fig 4C-D).

To compare the difference of filopodia dynamics between vehicle-treated infected cells and actin inhibitors treated infected cells, we performed the same quantification as in Figure 1 upon drug treatment with or without infection. Thereinto, VLP infection can still increase the length, number, protrusion speed and retraction speed of filopodia in blebbistatin-treated cells, but both parameters were significantly lower than the vehicle-treated cells (Fig 4E-I). Of note, VLP infection can also induce longer filopodia formation and increase the movement of filopodia, but cannot increase the dynamics of filopodia in SMIFH2-treated cells (Fig 4C, 4E-H). Even the formation and dynamics of filopodia were increased, VLP infection cannot change the movement speed of filopodia in CK666-treated cells (Fig 4E-H). Specifically, VLP infection cannot stimulate the dynamics, movement speed of filopodia, as well as longer filopodia formation in fascin inhibitor NP-G2-044 treated cells (Fig 4E-H). Compared with DMSO-treated cells, NP-G2-044 treatment kept the cells in a stationary state, which caused the reduction of VLP entry. Taken together, these results show that efficient entry of SARS-CoV-2 VLPs is dependent on intact actin filaments and filopodia dynamics.

### Defected host cell filopodia compromises the invasion of authentic SARS-CoV-2

To elaborate whether ‘surfing’ and ‘grabbing’ rely on actin filaments and filopodia components, we performed the same drug treatment and VLP infection as in Figure 4. Combining the results of three independent experiments, we found that drug treatments decreased the frequency of events in which the VLP particles entered the cell via filopodia (Fig 5A-E). Quantitative analysis revealed that NP-G2-044, SMIFH2 and CK666 treatment significantly decreased the real-time velocity of VLPs before entry, and in turn increase the VLPs entry time, except for blebbistatin (Fig 5 F-G). These experiments indicated that transport of VLPs on filopodia is actin-dependent, and is its components formin, fascin and Arp2/3 based.

**Figure 5.**
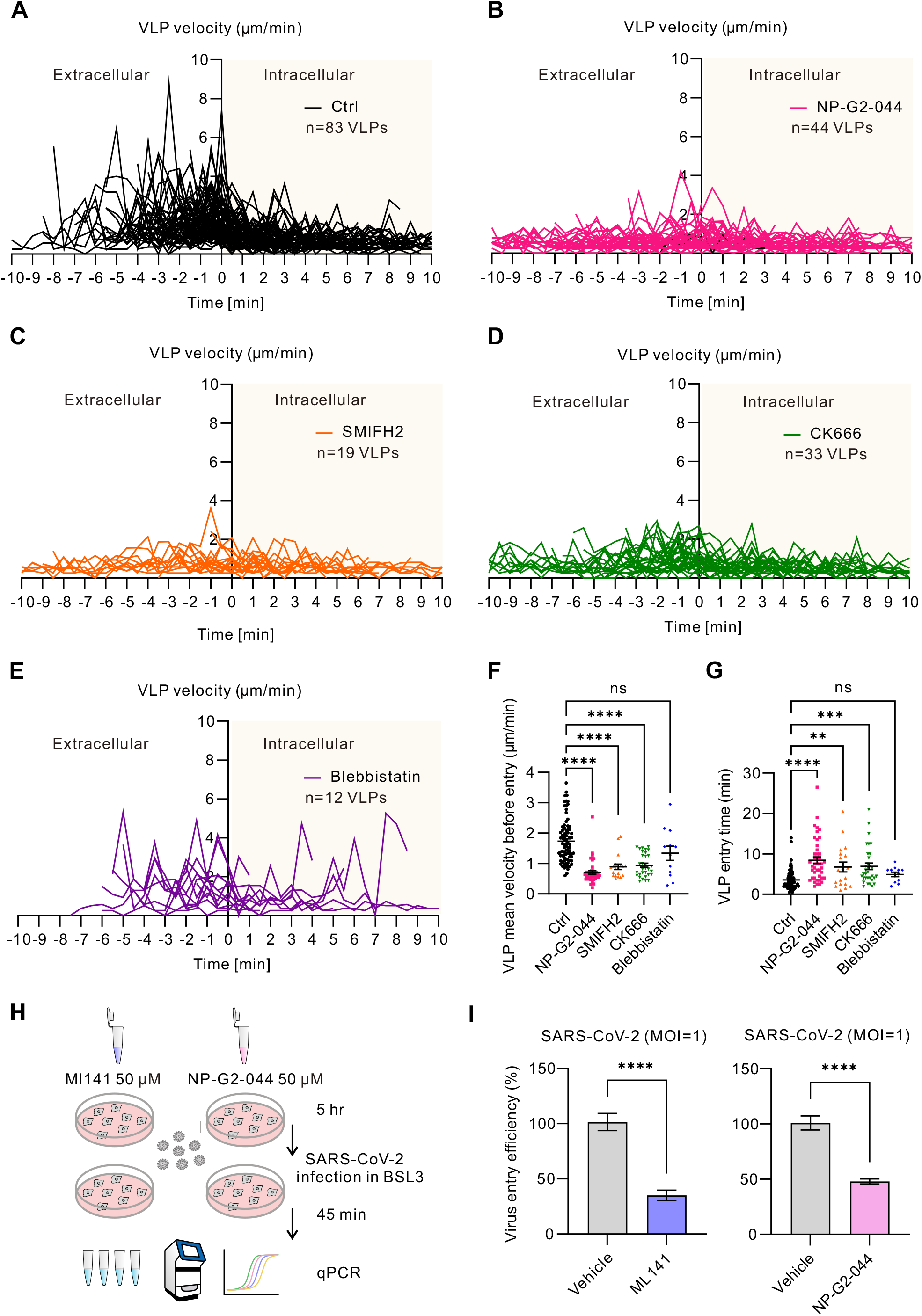
Quantification analysis of the effects of actin components inhibitors on SARS-CoV-2 entry. (**A-E**) The tracks of transient speed of VLP particles during entry from extracellular into intracellular cytoplasm upon untreated (Ctrl) (A), NP-G2-044 (50 μM for 2 h) (B), SIMFH2 (15 μM for 1 h) (C), CK666 (100 μM for 2 h) (D), Blebbistatin (17 μM for 1 h) (E), respectively. The original point on the X axis represents the viral entry time point. The negative number on the X axis represents the time before the virus enters the cell. The positive number on the X axis represents the time after the virus enters the cell. (**F**) Quantification of the mean velocities of VLP prior entering the untreated and inhibitors-treated cells. (**G**) Quantification of the duration of VLP particles enter into the untreated and inhibitors-treated cells. (**H**) Schematic diagram of drug treatment and authentic SARS-CoV-2 infection. (**I**) Viral entry efficiency of SARS-CoV-2 in BSL3. VeroE6-ACE2 cells were pretreated with 50 μM ML141 or 50 μM NP-G2-044 for 5 h, and challenged with SARS-CoV-2 (MOI=1) for 45 min. Viral RNA levels of internalized virus were measured by qRT-PCR. The data are normalized to the mean of DMSO-treated cells and represent as means ± SEM from three independent experiments. ns (no significant difference), *P* > 0.05, \**P* ≤ 0.05, \*\**P* ≤ 0.01, \*\*\**P* ≤ 0.001 and \*\*\*\**P* ≤ 0.0001 (One-way ANOVA test).

To further verify the suppressing effect of filopodia inhibitors on virus entry, we used authentic SARS-CoV-2 (WIV04 strain) and performed viral entry assay in BSL3 laboratory. In brief, VeroE6-ACE2 cells were pre-treated with Cdc42 inhibitor ML141 (50 μM for 5h) and fascin inhibitor NP-G2-044 (50 μM for 5h), respectively. Then, cells were infected with SARS-CoV-2 (MOI=1) for 45 min at 37°C, and treated with proteinase K to eradicate free virions. Total cellular RNA was extracted and the intracellular viral RNA level was determined by RT-qPCR (Fig 5H). The results showed that both pharmacological treatments can reduce the amount of viral entry by more than 50% (Fig 5I), which suggest the invasion of SARS-CoV-2 is indeed dependent on activated Cdc42 and functional fascin.

## Discussion

COVID-19 pandemic caused by SARS-CoV-2 poses a severe threat to global health due to the high transmissibility, broad cell tropism, and post-acute sequelae (Groff et al. 2021, Ramos da Silva et al. 2021, Lee et al. 2022). Our findings in this study highlight the crucial function of filopodia in the process of cell-free-virus invasion (Fig 6), which complement the previously reported role of filopodia during virus egress and cell-cell-transmission (Bouhaddou et al. 2020, Barreto-Vieira et al. 2021, Nijenhuis et al. 2021, Pepe et al. 2022, Swain et al. 2022).

**Figure 6.**
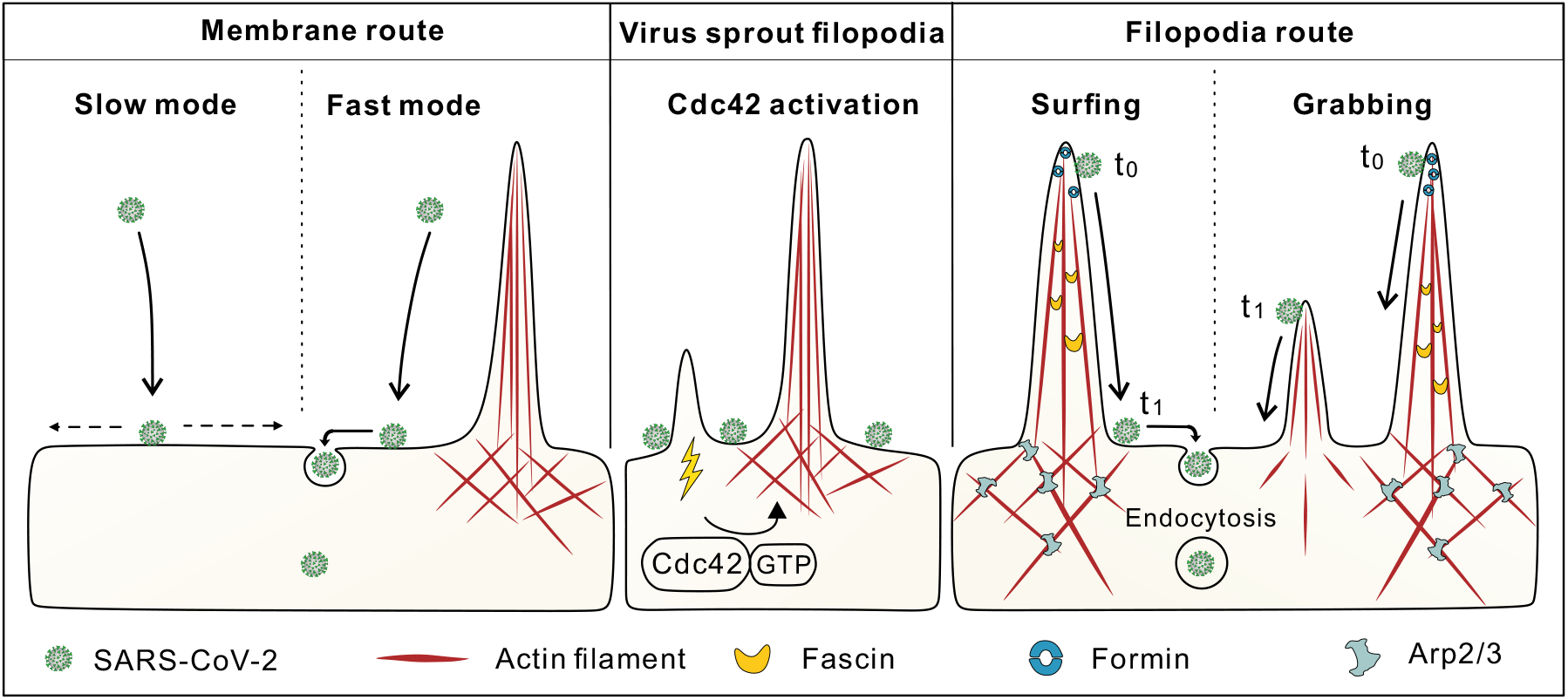
Schematic diagram of SARS-CoV-2 induced filopodia formation in host cells to facilitate viral entry.

Cell surface features, like filopodia, are essential for engaging and responding to pathogens. Numbers of pathogens have been shown to stimulate the formation of cellular filopodia (Horwitz 1984, Lee and Ng 2004, Rottner et al. 2005, Quetglas et al. 2012, Mehedi et al. 2017). Boosting cellular filopodia is a common strategy employed by pathogens to facilitate further invasion, and SARS-CoV-2 appears also take advantage of this efficient avenue based on our findings. Interestingly, our experiments reveal that the enhancement of filopodia in host cells appears to be irrelevant to the expression level of cellular ACE2 (Fig 1A,1C,S1G), nevertheless the entry efficiency of VLP particles is dependent on the level of ACE2 (Fig S1E), indicating that there are common mechanisms for virus-induced filopodia formation.

From the physical perspective, enhanced active stress can break the symmetry of the cell and stimulate filopodia formation (Fig 1H-I). From the biochemical perspective, Ras-related superfamily of small GTPases can trigger the formation of actin-based structures (Nobes and Hall 1995, Tapon and Hall 1997, Hall 1998). Correspondingly, numbers of viruses can control over the formation of actin-rich filopodia through modulating Rac, Rho, Cdc42, which in turn enhance virus uptake (Ohta et al. 1999, Puls et al. 1999, Smith et al. 2008, Zamudio-Meza et al. 2009, Nikolic et al. 2011, Taylor et al. 2011, Xiang et al. 2012, Freeman et al. 2014, Chang et al. 2022, Hunziker et al. 2022). Recent study reported that SARS-CoV-2 infection induces the upregulation of actin cytoskeleton via the kinase effectors of Rho GTPases (Liu et al. 2020). However, the relationship between Rho GTPases and increased filopodia in the entry of SARS-CoV-2 remain unclear. Our data first correlates the filopodia dynamics during SARS-CoV-2 VLP infection with the active level of Cdc42 (Fig 1J-O).

Different from recent reports that SARS-CoV-2 induced filopodia plays an important role in virus spreading (Bouhaddou et al. 2020, Barreto-Vieira et al. 2021, Nijenhuis et al. 2021, Pepe et al. 2022, Swain et al. 2022), we identified two filopodia-dependent particle entry processes ‘viral surfing’ and ‘viral grabbing’ that SARS-CoV-2 VLPs utilized for their invasion (Fig 2B-C). In surfing pattern, VLP particles directly trafficking along already-established filopodia to reach the membrane of host cells for entry, which is utilized by many viruses such as murine leukemia virus (MLV) (Lehmann et al. 2005), herpes simplex virus type 1 (HSV-1) (Dixit et al. 2008, Oh et al. 2010) and vaccinia virus (VV) (Mercer and Helenius 2008). However, ‘viral grabbing’ is rarely reported. To our knowledge, a similar action only occurs when filopodia of macrophages or epithelial cells retract and subsequently pull bound particle back to the cell (Kress et al. 2007, Mattila and Lappalainen 2008). Moreover, we found that the mean pulling velocity of VLP-loaded filopodia is substantially slower than unencumbered filopodia. According to simulation results, the difference in filopodial retraction velocities depend on the friction resistance at substrate surface (Fig 2H), that is, the counteracting forces produced by different loads as well as the density of loads (Cojoc et al. 2007, Kress et al. 2007, Michiels et al. 2022). The relationship between SARS-CoV-2 induced filopodial retraction force and viral entry efficiency remains to be characterized.

Compared with plasma membrane route, both of two patterns via filopodia enable VLP particles to enter the host cell quickly without any random forward-backward hesitation as we often observed on plasma membrane (Fig 2B-C,3A-F). ‘Surfing’ and ‘grabbing’ by filopodia may have evolved to avoid having to penetrate the microvilli-rich mucosal surfaces and the dense cortical actin cytoskeleton (Lehmann et al. 2005), which enhance the invasion of SARS-CoV-2. For several viruses other than SARS-CoV-2, association of virus and filopodia is not coincidental, but represent a common efficient infectious pathway that result in a direct transport of virus into endocytic hot spots (Helenius et al. 1980, Lehmann et al. 2005, Mercer and Helenius 2008, Schelhaas et al. 2008, Smith et al. 2008, Akhtar and Shukla 2009, Choudhary et al. 2013, Chang et al. 2016, Aliyu et al. 2021, Chang et al. 2022, Kloc et al. 2022). However, the underlying different mechanisms between these two strategies are worthy to be elaborated further.

Moreover, we tested the role of several important molecular compositions of filopodia in SARS-CoV-2 VLPs entry process. A similar phenomenon was found in exosomal uptake, which revealed that exosomes can efficiently and quickly enter primary human fibroblasts through surfing on filopodia, and loss of filopodia from the cell surface by formin inhibitor SMIFH2 treatment led to a reduction of exosome uptake (Schneider and Simons 2016). The similarities between exosomes and viruses suggest the commonality of the molecular mechanism of filopodia in sensing and delivery cargoes. Collectively, results in this study revealed that filopodia actively contribute to the invasion of emerging SARS-CoV-2, and these important filopodia component proteins might be new potential therapeutic targets for the treatment of COVID-19 disease.

## Material and Methods

### Cell culture

African green monkey kidney epithelial cells (VeroE6) cells and human embryonic kidney cell 293T (HEK293T) cells were maintained in high glucose Dulbecco’s modified Eagle’s medium (DMEM) (1055-57-1A, Biological Industries-1055-57-1A, Biological Industries) supplemented with 10% fetal bovine serum (FBS) (#10270-106, Gibco), 100 U/ml Penicillin and 100 μg/ml Streptomycin (#SV30010, Hyclone) at 37°C in humidified atmosphere with 5% CO_2_. To construct ACE2 stable overexpressing VeroE6 cell line, The human ACE2 coding sequence was amplified and inserted into the vector plasmid pLV-EF1α-IRES-Puro (#85132, Addgene) for transient expression in HEK293T cells to obtain virus containing the target gene. In brief, 2×10^5^ VeroE6 cells were co-transfected with 2 μg ACE2-overexpressing plasmid, together with 1 μg pVSV-G (#138479, Addgene) and 2 μg psPAX2 (#12260, Addgene) packaging plasmid by using polyethylenimine (Xiang et al. 2012) (#23966-1, Polysciences) reagent as manufacturers’ instruction. The culture supernatants were harvested at 48 h post-transfection and filtered through 0.45 μm filters. VeroE6 cells transduced with lentiviruses containing hACE2 were selected with 2 μg/ml puromycin to obtain ACE2-overexpressing VeroE6 cell clones.

### Production of SARS-CoV-2 VLPs

SARS-CoV-2 E, M and N genes (Wuhan-Hu-1, GenBank: QHD43419.1) were synthesized with optimization for Homo sapiens codon usage (Sangon Biotech and Tsingke Biotechnology) and cloned into pcDNA3.1 vector. The pcDNA3.1-eGFP-S plasmid was purchased from Genscript. We reconstructed pcDNA3.1 plasmids coding membrane, nucleocapsid and envelope proteins. The sequences of the genes above were optimized to have the highest frequency of codons in Homo sapiens and synthesized by Sangon Biotech and Tsingke Biotechnology. The above plasmids encoding viral structural proteins were co-transfected into HEK293T cells with an equimolar ratio using Lipofectamine 3000 (Invitrogen). The supernatant containing VLPs-EGFP were harvested 48 hours post transfection by centrifugation at 2000 × g for 15 minutes at 4°C. Clarified VLP-containing media was filtered through a 0.22 μm filter and then loaded on the top of a 20% sucrose cushion, followed by ultracentrifugation at 28,000 rpm for 4 hours at 4°C with a Beckman Type SW32 rotor. Finally, VLP-containing pellets were gently resuspended in the TNE buffer (10 mM Tris, 100 mM NaCl, 1 mM EDTA, pH 7.6) and dialyzed at 4°C in the dark. SARS-CoV-2 VLPs consisting of wild-type S protein were generated by a similar method.

### Transmission electron microscopy

Purified VLPs were loaded onto carbon-coated copper grids. The grids were washed once with PBS, and subsequently stained with 10 μl of 1% phosphotungstate and air dried. Samples were visualized under a transmission electron microscope (H-7000 FA, Hitachi).

### Treatments with chemical inhibitors and VLPs infection

In brief, 35 to 40% confluent monolayer of VeroE6-ACE2 cells was seeded in a 35 mm confocal dishes (Φ20mm glass) (#FCFC020, Beyotime), and incubated with 0.1 μM latrunculin B (#sc-203318, Santa Cruz) for 0.5 h, or 17 μM blebbistatin (#B0560; Sigma) for 1 h, or 50 μM NP-G2-044 (#S2962, Selleck) with for 2 h, or 100 μM CK666 (#sml0006, Sigma) for 2 h, or 10 μM SMIFH2 (#344092, EMD Millipore) for 1 h, or 50 μM ML141 (#HY-12755, MedChem Express) for 5 h, or 500 ng/mL Bradykinin (#TP1277, TargetMol) for 5 h, in 2% FBS-DMEM medium. DMSO (#A3672, PanReac AppliChem) was used as a control for latrunculin B, blebbistatin, NP-G2-044, CK666, SMIFH2 and ML141. Sterile ddH_2_O was used as a control for Bradykinin. After incubation, 0.15 mg VLPs were added to 1 mL drug-containing media and further incubated for 0.5 h or 3 h. Finally, cells were processed for live-cell imaging or immunofluorescence labeling.

### Cytotoxicity assay

1×10^4^ VeroE6-ACE2 cells were seeded in a 96-well plate and allowed to adhere and spread overnight. The commercially available CytoTox 96® Non-Radioactive Cytotoxicity Assay (#G1780, Promega) was performed after drug treatments as manufacturers’ instruction. In brief, 50 μL cell supernatant was transferred to a new 96-well plate, added 50 μL CytoTox 96® reagent and incubated for 30 min at room temperature (RT). The enzymatic reaction was stopped using 50 μL Stop Solution and then the absorbance at 490 nm was measured.

### Cdc42 activity assay

9×10^5^ VeroE6-ACE2 cells were seeded in a 6 cm dish, infected with 4.5 mg EGFP-VLP for 1 h at 37°C. For cell lysis collection, cells were washed with PBS, lysed with 700 μL ice-cold 1×Lysis/Binding/Wash Buffer plus protease and phosphatase inhibitors (Beyotime, #P1045), scraped, incubated on ice for 5 min, and then centrifuged at 6000×g for 15 min at 4°C. For affinity precipitation of activated G protein, 100 μL 50% resin slurry were added to the spin cup, centrifuged at 6000×g for 30 s, washed with 400 μL 1×Lysis/Binding/Wash Buffer, then incubated with 20 μg GST-PARK1-PBD and 700 μL cell lysate at 4°C on a shaker with low speed for 1 h. For elution, beads were spun down and washed with 400 μL 1×Lysis/Binding/Wash Buffer for three times, then incubated with 50 μL reducing sample buffer containing with 200 mM DTT at RT for 2 min, and centrifuged at 6000×g for 2 min. The elution samples were boiled for 5 min and subjected to SDS-PAGE, followed by western blotting.

### Immunofluorescence microscopy

Immunofluorescence (IF) experiments were performed as previously described (Jiu et al., 2019). Briefly, cells were fixed with 4% PFA in PBS for 15 min at RT, washed three times with PBS, and permeabilized with 0.1% Triton X-100 in PBS for 5 min. Cells were then blocked in PBS supplemented with 5% bovine serum albumin (BSA) (#A23088, ABCONE). Both primary and secondary antibodies were applied onto cells and incubated at RT for 2 h. Alexa-conjugated phalloidin was added together with secondary antibody solutions onto cells. Cells were mounted in Fluoromount-G reagent (SountherBiotech-01, SountherBiotech) and imaged using Olympus spinSR10 Ixplore spinning disk confocal microscope with UplanApo 60×/1.5 Oil objective (Olympus Corporation, Japan). The following antibodies were used in this study: anti-SARS-CoV-2 spike glycoprotein rabbit polyclonal antibody (dilution 1:1000; #ab272420, Abcam); anti-SARS-CoV-2 nucleocapsid protein rabbit polyclonal antibody (dilution 1:1000; #ab273167, Abcam); Alexa Fluor 488 goat anti-rabbit IgG (H+L) (dilution 1:1000; #A11008, Invitrogen); Alexa Fluor 555 phalloidin (dilution 1:500; #A34055, Invitrogen).

### Live-cell imaging

For live-cell imaging, 35 mm glass-bottomed dishes (MatTek Corporation) were coated with 10 μg/ml fibronectin (#F2006, Sigma-Aldrich) in PBS for 2 h at 37°C, washed with PBS twice before seeding of cells. 1.6×10^4^ VeroE6-ACE2 cells were seeded and infected with 1.5 mg VLP for 0.5 h at 37°C, then incubated with Alexa 555-wheat germ agglutinin (WGA) (2 μg/ml; #W32464, Thermo Fisher) or Alexa 647-wheat germ agglutinin (WGA) (2 μg/ml; #W32466, Thermo Fisher) for 10 min at 37°C to visualize the plasma membrane. Image series were acquired on an Olympus spinSR10 Ixplore spinning disk confocal microscope using a 60× U plan apochromat high resolution objective with NA=1.5, with time interval of 30 s for 1-2 h. All live cell imaging data were further analyzed by Imaris 9.2 (Bitplane) and ImageJ software.

### Single-particle tracking data processing

To reduce the large intensity differences between the dense membrane signal and the isolated filopodia, we first used a percentile-based normalization, which is defined for an image *x* as

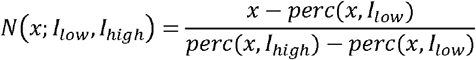

where *perc* (*x, I*) is the *I*-th percentile of all pixel values of *x*, and *I*_low_ and *I*_high_ represent the lowest and highest values, respectively. We used *I*_low_ as 0% for all images, and *I*_high_ as 99.8% and 100% for the WGA and VLP images, respectively. Then, all pixel values of images were affinely scaled to 0 and 1. Finally, the sparse deconvolution (Zhao et al. 2022) was used to eliminate the read-out noise and enhance the contrast of the low signal-to-noise ratio of images.

### Theoretical model

We developed an active gel model to capture the filopodial dynamics. A scalar field *ϕ* was introduced to describe the distribution of the material that makes up the cell. *ϕ* took value 1 in the cell and 0 otherwise. Cytoskeletal actin filaments exhibited directionality in the cell. We adopted the tensor **Q**=2*S*(**nn**-**I**/2) to describe the alignment of actin filaments, where *S* is the order parameter, **n** is the direction of individual filaments, and **I** denotes the two-dimensional unit tensor (Blow et al. 2017, Carenza et al. 2019). The free energy *(F)* of the system can be written as

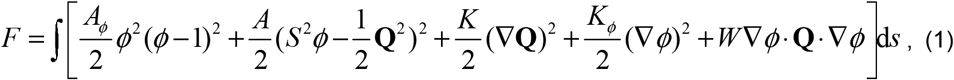

where *A_ϕ_* and *A* are material parameters, *K* is the elastic coefficient, *K_ϕ_* is the surface tension coefficient, and *W* is the anchoring strength at the cell boundary. The evolutions of actin flow **v**, *ϕ* and **Q** were governed by

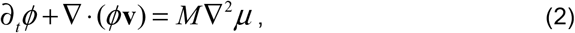

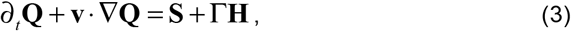

where *M* is the diffusion coefficient and the chemical potential is defined as *μ* = *δF / δϕ*. The tensor

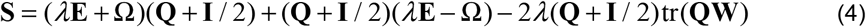

stands for the corotation term, which determines the alignment of cytoskeleton in response to gradients in the velocity field **v** (Blow et al. 2017, Li et al. 2020); **W** = ▽**v**, with **E** and **Ω** denoting the symmetrical and antisymmetric parts of **W**; *λ* is the alignment coefficient which controls the relative dominance of **E** and **Ω;** Γ is the rotational diffusivity, and **H** is the molecular field determined by the free energy *F*, that is, **H** = –*δF/ δ***Q**. Consider for simplicity the incompressible condition with constant density *ρ* The fluid velocity **v** satisfies **▽**·**v** = 0 (Blow et al. 2017). The governing equation of **v** was given as

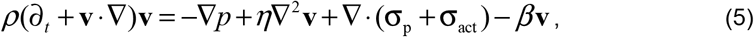

where *η* is the viscosity coefficient, *p* is hydrodynamic pressure, and *β* is friction coefficient characterizing the resistance from the underlying substrate. The passive stress can be expressed as

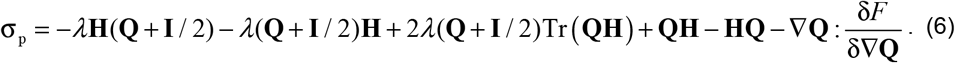

The active stress was taken as **σ**_act_ = -*ζ***Q**, which describes the extensile or contractile interaction of cytoskeleton due to cellular activity (Marchetti et al. 2013, Julicher et al. 2018, Li et al. 2020). We employed the hybrid lattice Boltzmann method to numerically solve equations (2), (3), and (5). Our simulation areas were taken as square windows with grids of 200 × 200. Two geometrical boundaries including circular and rectangular domains were considered to simulate the unconstraint protrusion of filopodia in a host cell and the growth of a single filopodium. For both domains, we set *ϕ*=1 inside the circle or rectangle. For rectangular domain, we allowed the diffusion only in the *y*-direction. To facilitate simulation, we used the simulated space and time steps as the space and time units for normalization. The typical parameter values were taken as *A*=0.1, *K*=0.02, *A_ϕ_* =0.2, *K_ϕ_* =0.05, *W*=0.05, *M*=0.1, Λ=0.9, Γ=0.1, and η=1/6.

### Authentic SARS-CoV-2 infection in BSL3 laboratory

VeroE6-ACE2 cells were pre-incubated with NP-G2-044 (50 μM), ML141 (50 μM) or DMSO solvent for 5 h. Cells were infected with authentic SARS-CoV-2 (WIV04) at a multiplicity of infection (MOI) of 1 for 45 min at 37°C. Cells were washed with PBS and then treated with proteinase K (Beyotime; final concentration, 1 mg/mL) for 30 min at 4°C to eradicate free virions. Then cells were washed with PBS and treated with TriZol® LS reagent (Invitrogen) to inactivate infectious viruses. Total RNA was extracted from infected cells using Direct-zol™ RNA MiniPrep Kits (Zymo). Viral RNA level was determined by real-time quantitative PCR (RT-qPCR) with human GAPDH as endogenous control. RT-qPCR was performed by using a HiScript II one-step qRT-PCR SYBR green kit (Vazyme) on a CFX96 real-time PCR detection system (Bio-Rad Laboratories, Inc.). The following primers were used: SARS-CoV-2-forward, 5’-CAATGGTTTAACAGGCACAGG-3’; SARS-CoV-2-reverse, 5’-CTCAAGTGTCTGTGGATCACG-3’ (Jiang RD et al. 2020); GAPDH-forward, 5’-GAAGGTGAAGGTCGGAGTC-3’; GAPDH-reverse, 5’-GAAGATGGTGATGGGGCTTC-3’. Results are expressed as the fold difference using expression in vehicle treated cells as calibrator value.

### Data analysis

The measurement of the parameters of filopodia is conducted in ImageJ. The measurement of VLP velocity is conducted in ImageJ with manual tracking plugin. The measurement of VLP number is conducted in Imaris with Spots tool. Statistical analyses were conducted using Graphpad Prism 9 (GraphPad) and significance was determined by unpaired Student’s *t* test, one-way ANOVA or two-way ANOVA. *p* values were indicated by \**P*<0.05, or \*\**P*<0.01, or \*\*\**P*<0.001, or \*\*\*\**P*<0.0001. All data were presented as mean ± SEM.

## Author Contributions

Y.Z. carried out the VLP infection experiments and interpretation of the data. X.Z. and H.Y. carried out VLP construction and the authentic virus infection experiments in BSL3 laboratory. B.L. and Z.L. performed the simulation. H.L. and W.Z. analyzed the single particle tracking data. D.T., S.Z. and Q.Z. participated in the data quantification. Y.J. and Z.C. supervised the whole project. Y.J. and Y.Z. conceived the study, and wrote the manuscript with contributions from P.L. and H.L. and all other authors.

## Acknowledgement

This study was supported by the Chinese Academy of Science-Vice Presidency Science and Technology Silk Road Science Fund (GJHZ2021138); Key Research and Development Program, Ministry of Science and Technology of China (2021YFC2300204); National Natural Science Foundation of China (32222022, 92054104, 31970660, 31925025); Strategic Priority Research Program of the Chinese Academy of Sciences (No. XDB29050201).

## References

1. Akhtar, J. and D. Shukla (2009). “Viral entry mechanisms: cellular and viral mediators of herpes simplex virus entry.” FEBS J 276(24): 7228–7236.

2. Aliyu, I. A., A. S. Kumurya, J. A. Bala, H. Yahaya and H. Saidu (2021). “Proteomes, kinases and signalling pathways in virus-induced filopodia, as potential antiviral therapeutics targets.” Rev Med Virol 31(5): 1–9.

3. Barreto-Vieira, D. F., M. A. N. da Silva, C. C. Garcia, M. D. Miranda, A. D. R. Matos, B. C. Caetano, P. C. Resende, F. C. Motta, M. M. Siqueira, W. Girard-Dias, et al. (2021). “Morphology and morphogenesis of SARS-CoV-2 in Vero-E6 cells.” Mem Inst Oswaldo Cruz 116: e200443.

4. Bayati, A., R. Kumar, V. Francis and P. S. McPherson (2021). “SARS-CoV-2 infects cells after viral entry via clathrin-mediated endocytosis.” J Biol Chem 296: 100306.

5. Blow, M. L., M. Aqil, B. Liebchen and D. Marenduzzo (2017). “Motility of active nematic films driven by “active anchoring”.” Soft Matter 13(36): 6137–6144.

6. Bouhaddou, M., D. Memon, B. Meyer, K. M. White, V. V. Rezelj, M. Correa Marrero, B. J. Polacco, J. E. Melnyk, S. Ulferts, R. M. Kaake, et al. (2020). “The Global Phosphorylation Landscape of SARS-CoV-2 Infection.” Cell 182(3): 685–712 e619.

7. Carenza, L. N., G. Gonnella, D. Marenduzzo and G. Negro (2019). “Rotation and propulsion in 3D active chiral droplets.” Proc Natl Acad Sci U S A 116(44): 22065–22070.

8. Chang, K., J. Baginski, S. F. Hassan, M. Volin, D. Shukla and V. Tiwari (2016). “Filopodia and Viruses: An Analysis of Membrane Processes in Entry Mechanisms.” Front Microbiol 7: 300.

9. Chang, K., H. Majmudar, R. Tandon, M. V. Volin and V. Tiwari (2022). “Induction of Filopodia During Cytomegalovirus Entry Into Human Iris Stromal Cells.” Front Microbiol 13: 834927.

10. Choudhary, S., L. Burnham, J. M. Thompson, D. Shukla and V. Tiwari (2013). “Role of Filopodia in HSV-1 Entry into Zebrafish 3-O-Sulfotransferase-3-Expressing Cells.” Open Virol J 7: 41–48.

11. Cojoc, D., F. Difato, E. Ferrari, R. B. Shahapure, J. Laishram, M. Righi, E. M. Di Fabrizio and V. Torre (2007). “Properties of the force exerted by filopodia and lamellipodia and the involvement of cytoskeletal components.” PLoS One 2(10): e1072.

12. Cudmore, S., P. Cossart, G. Griffiths and M. Way (1995). “Actin-based motility of vaccinia virus.” Nature 378(6557): 636–638.

13. Dixit, R., V. Tiwari and D. Shukla (2008). “Herpes simplex virus type 1 induces filopodia in differentiated P19 neural cells to facilitate viral spread.” Neurosci Lett 440(2): 113–118.

14. Freeman, M. C., C. T. Peek, M. M. Becker, E. C. Smith and M. R. Denison (2014). “Coronaviruses induce entry-independent, continuous macropinocytosis.” mBio 5(4): e01340-01314.

15. Giomi, L., M. J. Bowick, X. Ma and M. C. Marchetti (2013). “Defect annihilation and proliferation in active nematics.” Phys Rev Lett 110(22): 228101.

16. Groff, D., A. Sun, A. E. Ssentongo, D. M. Ba, N. Parsons, G. R. Poudel, A. Lekoubou, J. S. Oh, J. E. Ericson, P. Ssentongo, et al. (2021). “Short-term and Long-term Rates of Postacute Sequelae of SARS-CoV-2 Infection: A Systematic Review.” JAMA Netw Open 4(10): e2128568.

17. Hall, A. (1998). “Rho GTPases and the actin cytoskeleton.” Science 279(5350): 509–514.

18. Helenius, A., J. Kartenbeck, K. Simons and E. Fries (1980). “On the entry of Semliki forest virus into BHK-21 cells.” J Cell Biol 84(2): 404–420.

19. Horwitz, M. A. (1984). “Phagocytosis of the Legionnaires’ disease bacterium (Legionella pneumophila) occurs by a novel mechanism: engulfment within a pseudopod coil.” Cell 36(1): 27–33.

20. Hunziker, A., I. Glas, M. O. Pohl and S. Stertz (2022). “Phosphoproteomic profiling of influenza virus entry reveals infection-triggered filopodia induction counteracted by dynamic cortactin phosphorylation.” Cell Rep 38(4): 110306.

21. Jackson, C. B., M. Farzan, B. Chen and H. Choe (2022). “Mechanisms of SARS-CoV-2 entry into cells.” Nat Rev Mol Cell Biol 23(1): 3–20.

22. Julicher, F., S. W. Grill and G. Salbreux (2018). “Hydrodynamic theory of active matter.” Rep Prog Phys 81(7): 076601.

23. Kloc, M., A. Uosef, J. Wosik, J. Z. Kubiak and R. M. Ghobrial (2022). “Virus interactions with the actin cytoskeleton-what we know and do not know about SARS-CoV-2.” Arch Virol 167(3): 737–749.

24. Koch, J., Z. M. Uckeley, P. Doldan, M. Stanifer, S. Boulant and P. Y. Lozach (2021). “TMPRSS2 expression dictates the entry route used by SARS-CoV-2 to infect host cells.” EMBO J 40(16): e107821.

25. Kress, H., E. H. Stelzer, D. Holzer, F. Buss, G. Griffiths and A. Rohrbach (2007). “Filopodia act as phagocytic tentacles and pull with discrete steps and a load-dependent velocity.” Proc Natl Acad Sci U S A 104(28): 11633–11638.

26. Li, Z. Y., D. Q. Zhang, S. Z. Lin and B. Li (2020). “Pattern Formation and Defect Ordering in Active Chiral Nematics.” Phys Rev Lett 125(9): 098002.

27. Lee, J. W. and M. L. Ng (2004). “A nano-view of West Nile virus-induced cellular changes during infection.” J Nanobiotechnology 2(1): 6.

28. Lee, L. Y. W., S. Rozmanowski, M. Pang, A. Charlett, C. Anderson, G. J. Hughes, M. Barnard, L. Peto, R. Vipond, A. Sienkiewicz, et al. (2022). “Severe Acute Respiratory Syndrome Coronavirus 2 (SARS-CoV-2) Infectivity by Viral Load, S Gene Variants and Demographic Factors, and the Utility of Lateral Flow Devices to Prevent Transmission.” Clin Infect Dis 74(3): 407–415.

29. Lehmann, M. J., N. M. Sherer, C. B. Marks, M. Pypaert and W. Mothes (2005). “Actin-and myosin-driven movement of viruses along filopodia precedes their entry into cells.” J Cell Biol 170(2): 317–325.

30. Liu, H. L., I. J. Yeh, N. N. Phan, Y. H. Wu, M. C. Yen, J. H. Hung, C. C. Chiao, C. F. Chen, Z. Sun, J. Z. Jiang, et al. (2020). “Gene signatures of SARS-CoV/SARS-CoV-2-infected ferret lungs in short-and long-term models.” Infect Genet Evol 85: 104438.

31. Mallavarapu, A. and T. Mitchison (1999). “Regulated actin cytoskeleton assembly at filopodium tips controls their extension and retraction.” J Cell Biol 146(5): 1097–1106.

32. Marchetti, M. C., J. F. Joanny, S. Ramaswamy, T. B. Liverpool, J. Prost, M. Rao and R. A. Simha (2013). “Hydrodynamics of soft active matter.” Reviews of Modern Physics 85(3): 1143–1189.

33. Mattila, P. K. and P. Lappalainen (2008). “Filopodia: molecular architecture and cellular functions.” Nat Rev Mol Cell Biol 9(6): 446–454.

34. Mehedi, M., P. L. Collins and U. J. Buchholz (2017). “A novel host factor for human respiratory syncytial virus.” Commun Integr Biol 10(3): e1319025.

35. Mellor, H. (2010). “The role of formins in filopodia formation.” Biochim Biophys Acta 1803(2): 191–200.

36. Mercer, J. and A. Helenius (2008). “Vaccinia virus uses macropinocytosis and apoptotic mimicry to enter host cells.” Science 320(5875): 531–535.

37. Michiels, R., N. Gensch, B. Erhard and A. Rohrbach (2022). “Pulling, failing, and adaptive mechanotransduction of macrophage filopodia.” Biophys J.

38. Nijenhuis, W., H. G. J. Damstra, E. J. van Grinsven, M. K. Iwanski, P. Praest, Z. E. Soltani, M. M. P. van Grinsven, J. E. Brunsveld, T. de Kort, L. W. Rodenburg, et al. (2021). “Optical nanoscopy reveals SARS-CoV-2-induced remodeling of human airway cells.” 2021.2008.2005.455126.

39. Nikolic, D. S., M. Lehmann, R. Felts, E. Garcia, F. P. Blanchet, S. Subramaniam and V. Piguet (2011). “HIV-1 activates Cdc42 and induces membrane extensions in immature dendritic cells to facilitate cell-to-cell virus propagation.” Blood 118(18): 4841–4852.

40. Nobes, C. D. and A. Hall (1995). “Rho, rac, and cdc42 GTPases regulate the assembly of multimolecular focal complexes associated with actin stress fibers, lamellipodia, and filopodia.” Cell 81(1): 53–62.

41. Oh, M. J., J. Akhtar, P. Desai and D. Shukla (2010). “A role for heparan sulfate in viral surfing.” Biochem Biophys Res Commun 391(1): 176–181.

42. Ohta, Y, N. Suzuki, S. Nakamura, J. H. Hartwig and T. P. Stossel (1999). “The small GTPase RalA targets filamin to induce filopodia.” Proc Natl Acad Sci USA 96(5): 2122–2128.

43. Onyeaka, H., C. K. Anumudu, Z. T. Al-Sharify, E. Egele-Godswill and P. Mbaegbu (2021). “COVID-19 pandemic: A review of the global lockdown and its far-reaching effects.” Sci Prog 104(2): 368504211019854.

44. Pepe, A., S. Pietropaoli, M. Vos, G. Barba-Spaeth and C. Zurzolo (2022). “Tunneling nanotubes provide a route for SARS-CoV-2 spreading.” Sci Adv 8(29): eabo0171.

45. Puls, A., A. G. Eliopoulos, C. D. Nobes, T. Bridges, L. S. Young and A. Hall (1999). “Activation of the small GTPase Cdc42 by the inflammatory cytokines TNF(alpha) and IL-1, and by the Epstein-Barr virus transforming protein LMP1.” J Cell Sci 112 (Pt 17): 2983–2992.

46. Quetglas, J. I., B. Hernaez, I. Galindo, R. Munoz-Moreno, M. A. Cuesta-Geijo and C. Alonso (2012). “Small rho GTPases and cholesterol biosynthetic pathway intermediates in African swine fever virus infection.” J Virol 86(3): 1758–1767.

47. Ramos da Silva, S., E. Ju, W. Meng, A. E. Paniz Mondolfi, S. Dacic, A. Green, C. Bryce, Z. Grimes, M. Fowkes, E. M. Sordillo, et al. (2021). “Broad Severe Acute Respiratory Syndrome Coronavirus 2 Cell Tropism and Immunopathology in Lung Tissues From Fatal Coronavirus Disease 2019.” J Infect Dis 223(11): 1842–1854.

48. Ridley, A. J. (2006). “Rho GTPases and actin dynamics in membrane protrusions and vesicle trafficking.” Trends Cell Biol 16(10): 522–529.

49. Rottner, K., T. E. Stradal and J. Wehland (2005). “Bacteria-host-cell interactions at the plasma membrane: stories on actin cytoskeleton subversion.” Dev Cell 9(1): 3–17.

50. Schelhaas, M., H. Ewers, M. L. Rajamaki, P. M. Day, J. T. Schiller and A. Helenius (2008). “Human papillomavirus type 16 entry: retrograde cell surface transport along actin-rich protrusions.” PLoS Pathog 4(9): e1000148.

51. Schneider, A. and M. Simons (2016). “Catching filopodia: Exosomes surf on fast highways to enter cells.” J Cell Biol 213(2): 143–145.

52. Smith, J. L., D. S. Lidke and M. A. Ozbun (2008). “Virus activated filopodia promote human papillomavirus type 31 uptake from the extracellular matrix.” Virology 381(1): 16–21.

53. Swain, J., P. Merida, K. Rubio, D. Bracquemond, I. Aguilar-Ordoñez, S. Günther, G. Barreto and D. Muriaux (2022). “Reorganization of F-actin nanostructures is required for the late phases of SARS-CoV-2 replication in pulmonary cells.” 2022.2003.2008.483451.

54. Tapon, N. and A. Hall (1997). “Rho, Rac and Cdc42 GTPases regulate the organization of the actin cytoskeleton.” Curr Opin Cell Biol 9(1): 86–92.

55. Taylor, M. P., O. O. Koyuncu and L. W. Enquist (2011). “Subversion of the actin cytoskeleton during viral infection.” Nat Rev Microbiol 9(6): 427–439.

56. Twarock, S., M. I. Tammi, R. C. Savani and J. W. Fischer (2010). “Hyaluronan stabilizes focal adhesions, filopodia, and the proliferative phenotype in esophageal squamous carcinoma cells.” J Biol Chem 285(30): 23276–23284.

57. Xiang, Y., K. Zheng, H. Ju, S. Wang, Y. Pei, W. Ding, Z. Chen, Q. Wang, X. Qiu, M. Zhong, et al. (2012). “Cofilin 1-mediated biphasic F-actin dynamics of neuronal cells affect herpes simplex virus 1 infection and replication.” J Virol 86(16): 8440–8451.

58. Zamudio-Meza, H., A. Castillo-Alvarez, C. Gonzalez-Bonilla and I. Meza (2009). “Cross-talk between Rac1 and Cdc42 GTPases regulates formation of filopodia required for dengue virus type-2 entry into HMEC-1 cells.” J Gen Virol 90(Pt 12): 2902–2911.

59. Zhao, W., S. Zhao, L. Li, X. Huang, S. Xing, Y. Zhang, G. Qiu, Z. Han, Y. Shang, D. E. Sun, et al. (2022). “Sparse deconvolution improves the resolution of live-cell super-resolution fluorescence microscopy.” Nat Biotechnol 40(4): 606–617.

