## Supplemental Figure Legends for "SARS-CoV-2 infected cells sprout actin-rich filopodia that facilitate viral invasion"

**Figure S1 Construction and characterization of EGFP tagged SARS-CoV-2 VLP.** (**A**) Schematic representation of SARS-CoV-2 VLPs-EGFP constructions in mammalian expression system. (**B**) Electron microscopy of SARS-CoV-2 VLPs and SARS-CoV-2 VLPs-EGFP. Scale bar 100 nm. (**C**) Immunofluorescence assay of SARS-CoV-2 VLPs-EGFP using anti-S or anti-N antibody on coverslips. Scale bar 2 µm. (**D**) Western blot verified ACE2 overexpression in VeroE6 cells. GAPDH is used to verify equal sample loading. (**E**) Entry efficiency of SARS-CoV-2 VLPs-EGFP in VeroE6 cells and VeroE6-ACE2 cells. Scale bar 15 µm. (**F**) Quantification of the numbers of VLP per cell at different hours post-infection (hpi). (**G**) SARS-CoV-2 VLPs infection induces the filopodia formation in VeroE6 cells. Scale bar 15 µm.

**Figure S2 Cytotoxicity verification of used chemical inhibitors.** VeroE6-ACE2 cells were treated with 0.1 μM Latrunculin B for 40 min (**A**), 17 μM Blebbistatin for 1 h (**B**), 15 μM SMIFH2 for 1 h (**C**), 500 ng/mL bradykinin for 5 h (**D**), 50 μM ML141for 5 h (**E**), 50 μM NP-G2-044 for 2 h (**F**). The data are represented as Means ± SEM from four replicates. ns *P* > 0.05, ***P* ≤ 0.01 (unpaired *t*-test). The data of maximum LDH release are from the group of adding lysis buffer.

**Video S1 related to Figure 1A.** Time-lapse video of Vero-E6-ACE2 overexpressing cells in normal culture condition. The display rate is 25 frame per second. Scale bar 10 μm.

**Video S2 related to Figure 1A.** Time-lapse video of Vero-E6-ACE2 overexpressing cells infected by SARS-CoV-2 VLP-EGFP. The display rate is 25 frame per second. Scale bar 10 μm.

**Video S3 related to Figure 2B.** Time-lapse video of SARS-CoV-2 VLP-EGFP particle surfs on the filopodia of Vero-E6-ACE2 overexpressing cells. The display rate is 8 frame per second. Scale bar 2 μm.

**Video S4 related to Figure 2C.** Time-lapse video of SARS-CoV-2 VLP-EGFP particle grabbed by the filopodia of Vero-E6-ACE2 overexpressing cells. The display rate is 5 frame per second. Scale bar 2 μm.

**Video S5 related to Figure 3A.** Time-lapse video of SARS-CoV-2 VLP-EGFP particle enters Vero-E6-ACE2 overexpressing cells via plasma membrane in fast mode. The display rate is 10 frame per second. Scale bar 2 μm.

**Video S6 related to Figure 3B.** Time-lapse video of SARS-CoV-2 VLP-EGFP particle enters Vero-E6-ACE2 overexpressing cells via plasma membrane in slow mode. The display rate is 20 frame per second. Scale bar 2 μm.
