## Supplementary figures and images for "SARS-CoV-2 infected cells sprout actin-rich filopodia that facilitate viral invasion"

### Supplemental Figure 1-2

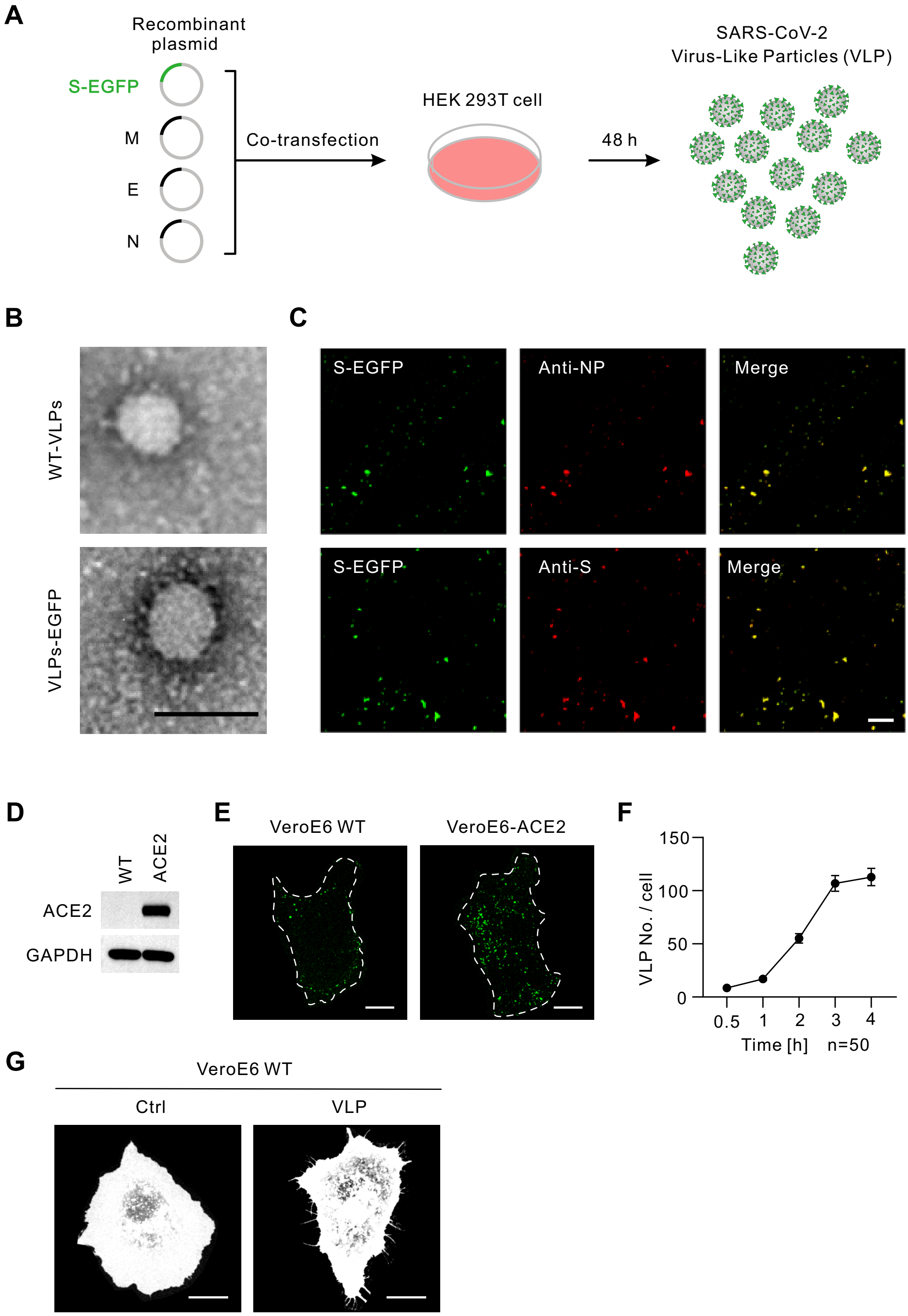

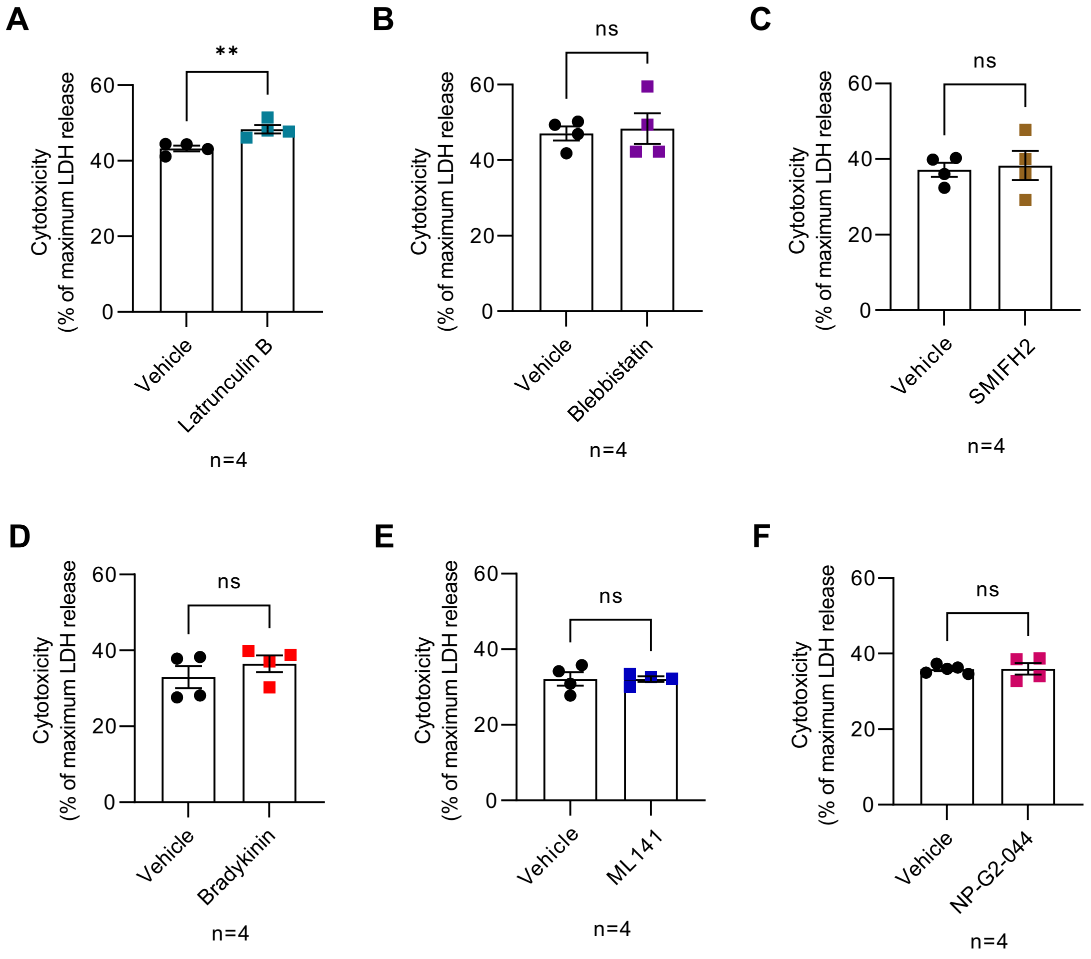
